# AI-based decoding of long covid cognitive impairments in mice using automated behavioral system and comparative transcriptomic analysis

**DOI:** 10.1101/2025.05.14.654036

**Authors:** Heba M. Amer, Mohamed M. Shamseldin, Sarah Faber, Mostafa Eltobgy, Amy Webb, Rabab El-Mergawy, Michelle Chamblee, Richard Perez, Owen Whitham, Asmaa Badr, Gauruv Gupta, Jihad Omran, Destiny Bissel, Jacob Yount, Estelle Cormet-Boyaka, Prosper N. Boyaka, Jianrong Li, Xiaoli Zhang, Mark E. Peeples, KC Mahesh, Maciej Pietrzak, Stephanie Seveau, Olga Kokiko-Cochran, Magdi Amer, Ruth M. Barrientos, Andrew Schamess, Eugene Oltz, Amal O. Amer

## Abstract

Long COVID (LC) following SARS-CoV-2 infection affects millions of individuals world-wide and manifests with a variety of symptoms including cognitive dysfunction also known as “brain fog”. This is characterized by difficulties in executive functions, planning, decision-making, working memory, impairments in complex attention, loss of ability to learn new skills and perform sophisticated brain tasks. No effective treatment options currently exist for LC-related cognitive dysfunction. Here, we use the IntelliCage, which is an automated tracking system of cognitive functions, following SARS-CoV-2 infection in mice, measuring the ability of each mouse within a group to perform tasks that mimic complex human behaviors, such as planning, decision-making, cognitive flexibility, and working memory. Artificial intelligence and machine learning analyses of the tracking data classified LC mice into distinct behavioral categories from non-infected control mice, permitting precise identification and quantification of complex cognitive dysfunction in a controlled, replicable manner. Importantly, we find that brains from LC mice with cognitive dysfunction exhibit transcriptomic alterations similar to those observed in humans suffering from LC-related cognitive impairments, including altered expression of genes involved in learning, executive functions, synaptic functions, neurotransmitters and memory. Together, our findings establish a validated murine model and an automated unbiased approach to study LC-related cognitive dysfunction for the first time, and providing a valuable tool for screening potential treatments and therapeutic interventions.

## Introduction

Post-acute COVID-19 syndrome (PACS) or Long COVID (LC) is characterized by persistent symptoms following acute SARS-CoV-2 infection, which last for more than 12 weeks, and cannot be explained by an alternative diagnosis ^1–5^. Studies have reported a high prevalence of cognitive decline in LC patients^6 7–13^, including functional deficits in executive function, memory, attention, inhibitory control, and processing speed that affect planning, decision-making, and problem-solving abilities ^10,14–19^. Neurological manifestations after SARS-CoV-2 infection are the leading contributor of disease burden and disability among all people suffering from LC ^20,21^. Therefore, there is an urgent need to better understand the pathobiology of LC and to enable the identification of drug targets as therapeutic candidates to manage LC-related cognitive dysfunction ^13^. However, clinical trials are difficult to plan and prioritize without better understanding of brain pathology, and without preclinical screening in animal models ^22^. A further hindrance is the lack of reliable behavioral testing techniques that can distinguish subtle differences and can be monitored for minor improvements in validated mice models of cognitive dysfunction post-SARS-CoV-2 infection.

Traditional behavioral assays, such as the Morris Water Maze ^23–25^, open field tests, and elevated mazes ^26–28^, have been widely used to assess cognitive behavioral function in neurodegenerative conditions, offering insights into spatial learning, memory, and exploratory drive. However, these approaches have several limitations. They primarily evaluate basic cognitive processes, cannot accurately assess higher-order cognitive deficits, may require prolonged testing periods, and are often performed in isolated conditions that induce stress ^24,29^. Furthermore, it can be difficult to track and interpret over time with repeated testing and within subject performance. Importantly, these techniques require handling, training and testing that can differ considerably across laboratories due to human variability ^30^.

While several murine models have been developed to investigate the chronic histological and pathological changes associated with LC, models specifically addressing LC-related cognitive dysfunction are lacking ^31,32^. This challenge is compounded by the fact that brain-related LC symptoms in humans are complex and not easily detected or quantified in mice by traditional methods. Thus, identification of diagnostic markers and therapeutic targets is hampered by the lack of LC model systems, thereby complicating prioritization of drug candidates to be used in clinical trials for LC-related cognitive dysfunction. An automated cage tracking system for mice, termed IntelliCage, provides unprecedented opportunity for longitudinal analysis of complex cognitive functions, enabling continuous monitoring and tracking of individual mice without direct human-animal interaction for over several weeks.

The system contains four operant testing corners accessible via tunnels, each equipped with two doors regulating access to water bottles and three LED indicators that can be used during cognitive testing ^33^. Each mouse to be placed in the IntelliCage system receives a subcutaneously injected RFID tag (transponder) several days prior to testing, which enables continuous monitoring and tracking of individual behaviors through ring antennas located in the corners of the cage without direct human-animal interaction ^33^. Mice interact with the system by performing nose-pokes to open the doors and access the reward (water), while the system continuously records their visits, nose-pokes, and drinking behavior. This setup allows for highly reproducible, standardized testing conditions, minimizing stress during multi-step task execution ^33,34^ and enhancing reliability ^35–38^. The automated cage system enables assessment of executive functions relevant to LC, including working memory, cognitive flexibility, response inhibition ^39^, learning speed, and adaptability ^30,40^. Experimental parameters can be precisely controlled, allowing mice to learn, relearn, and remember specific conditions to obtain rewards. Since LC patients frequently report deficits in problem-solving, executive function ^1,2^, attention, and processing speed ^41^, relevant IntelliCage-based tests were chosen and designed to mirror and test these impairments in mice.

Using this technology and a machine learning (ML) approach to data analysis, we report that LC mice display impaired executive functions and were less capable of performing sophisticated behaviors when compared to their uninfected counterparts. Secondary analysis of publicly available RNA sequencing (RNAseq) data of brain tissue obtained from deceased patients with LC-related cognitive dysfunction shows that the expression of genes involved in neuroinflammation, cognition, learning, neurotransmitters and memory pathways are significantly altered. To further investigate molecular correlates of LC-related cognitive dysfunction in mice, we performed RNAseq analysis of whole brain tissue in LC mice. Consistent with human data, significant transcriptional changes were observed in genes associated with pathways similar to those altered in human subjects with LC-related brain fog.

Together, we characterized an innovative mouse model of LC-related cognitive dysfunction using the IntelliCage system for the first time, which accurately detects behavioral changes weeks after SARS-CoV-2 infection. Cognitive deficits were supported by transcriptomic data linking altered molecular expression to executive functions and memory in humans and mice. This model advances our understanding of LC pathobiology and serves as a platform for studying cognitive dysfunction, identifying biomarkers and testing potential therapies.

## Results

### Working memory is altered in LC mice

Working memory is a limited capacity cognitive system that holds information temporarily to be utilized for reasoning, learning, decision-making and behavior ^39^. To study working memory in the LC model, mice were infected with SARS-CoV-2 MA10. Infected animals lost weight, then recovered gradually over two weeks (**Figure 1A** and **B**). The virus was cleared from the lungs by day 7 post-infection and no live virus was detected in the brain at any stage (**Figure 1C**). On day 55 post-infection, the LC and non-infected mice (control group) were injected with an RFID subcutaneously and then placed in the IntelliCage (**Figure 1A**). Mice were allowed to acclimatize to the cage for 5 days prior to testing. One day before the first test commenced, mice were conditioned to associate light cues from LED lights with access to water. This was done by programming the system to allow water access in all corners of the IntelliCage via a nose poke. Simultaneously, throughout the entirety of the drinking period, the three LED lights located in the corner would be on and would turn off once the mouse was done drinking. After the one-day conditioning period, the patrol and patrol reversal tests proceeded. During these tests, the “correct” corner-chamber (i.e., the corner where the mice could access water) was changed in a clockwise and counterclockwise manner, respectively, and when the mouse enters the correct corner, the LED lights above the doors would turn on ^42^, indicating to the mouse that they are in the “correct” corner. If the mouse went to an “incorrect” corner, the LED lights would remain inactive and should cue the mouse that they will not receive access to water.

**Figure 1:**
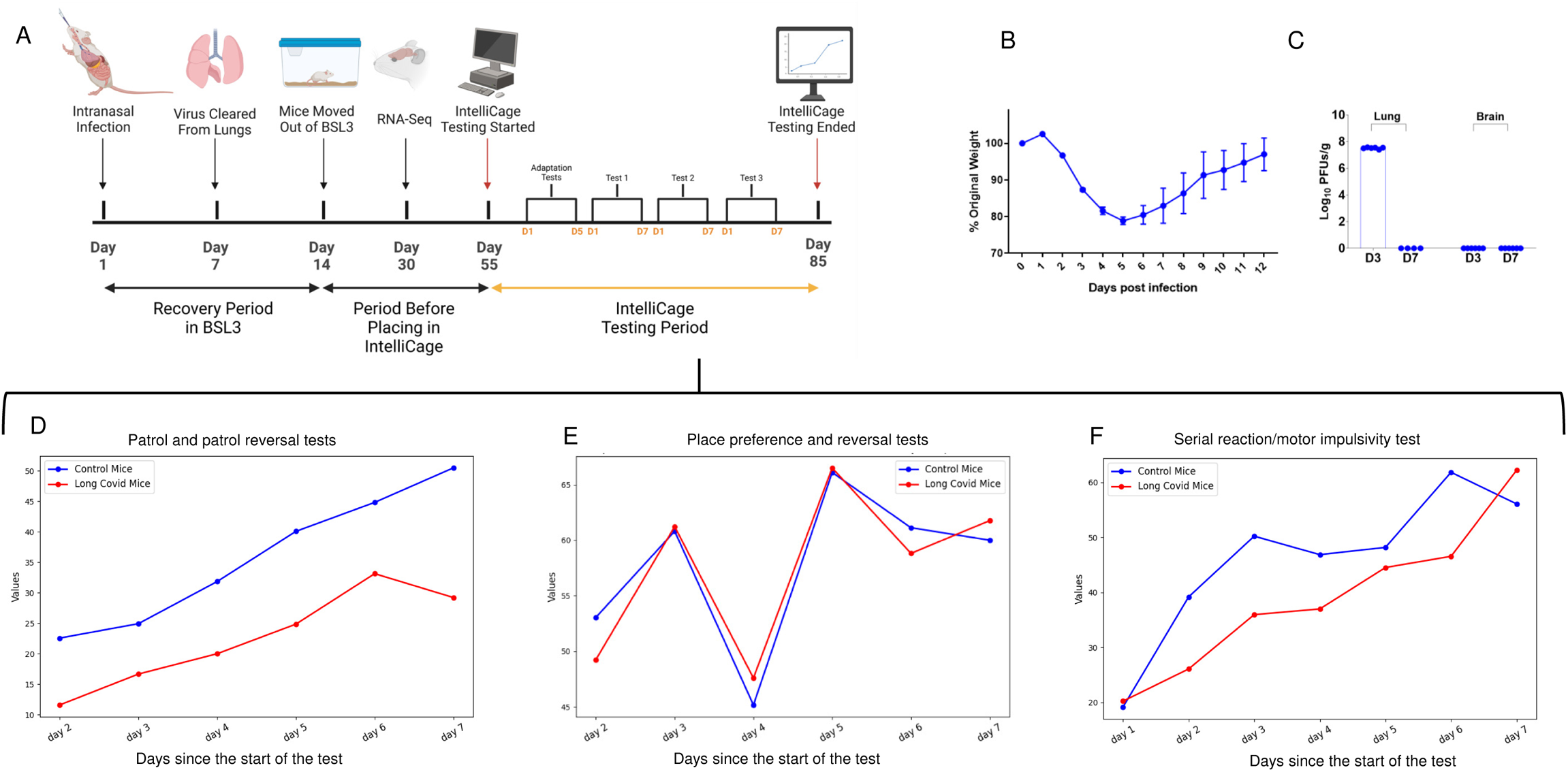
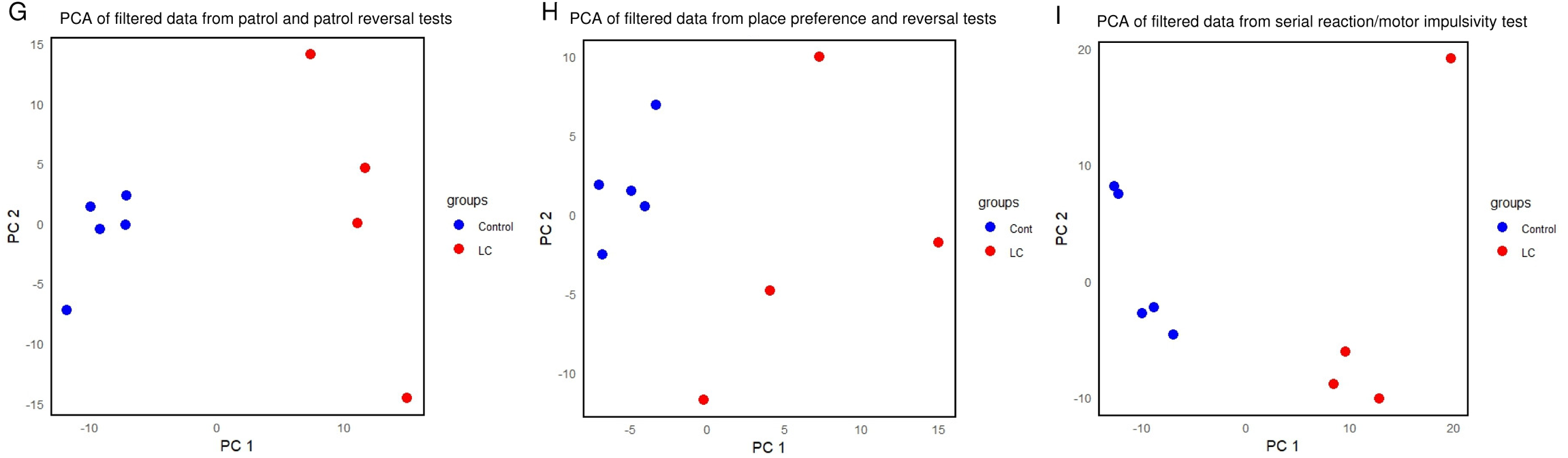
LC mice performance of complex behavior tests in the IntelliCage: **A**. Timeline (Created in BioRender. Amer, H. (2025) https://BioRender.com/0bumgen). **B**. Body weight loss of mice infected with SARS-CoV-2 displayed as a percentage of original weight at the start of the experiment. **C**. Viral titers expressed as log plaque forming units (PFUs) per gram of tissue, was quantified from lungs and brains of infected mice at 3- and 7-days post-infection. **D**. **Patrol and patrol reversal test.** Success rates of each group per day in correctly associating a lack of the positive cue with the inability to access water. The Y axis represents number of times a mouse entered an incorrect corner and did not perform a nose-poke when the lights did not go on out of the total incorrect visits. Control group (blue) achieved a significantly higher hourly total correct percentage (34.57%) when compared to the LC group (red) (21.70%) (p-value=8.15E-10). **E**. **Place preference/place learning.** The Y axis represents the percentage of correct visits with nose-pokes out of the total number of visits with nose pokes and was calculated per day. The overall correct percentage of the control group (blue) was 56.98% while the LC group (red) was 56.83% (p-value=0.08). **F**. **Higher-order cognitive function, and cognitive flexibility and impulsivity test.** The Y axis represents the percentage of successful trials out of total trials initiated (with elimination of semi-successful and semi-failed trials) per day in both groups. Control group (blue) (39.44%), LC group (31.97%) (p-value= 0.004). (**B-I**) (N=4) two sample t-test. (**G-I**) Unsupervised dimensionality reduction technique, Principal Component Analysis (PCA) on the filtered dataset of each test. Each point represents a data sample, projected onto the first two principal components (PC1 and PC2). The colors indicate different groups within the dataset. Blue represents the control uninfected mice and red represents the LC mice with cognitive dysfunction.

The patrol and patrol reversal tests allow evaluation of the animal’s working memory, their memory reference and their ability to process positive visual cues and take actions accordingly ^34,42^. The first parameter we measured was correct visits with nose-pokes out of total visits with nose-pokes. For this, there was no statistical significance between the LC (40.11%) when compared to the control group (39.24%) (P-value =0.28, two sample t-test) (**Figure 1D**). However, when measuring the ability of a mouse to connect the lack of the positive cue with the inability to access water – the number of times it entered an incorrect corner and did not perform a nose-poke when the lights remained off out of total incorrect visits – we found that the control group achieved a significantly higher hourly total correct response (34.57% no nosepokes) when compared to the LC group (21.70%) (p-value=8.15E-10, two sample t-test) (**Figure 1D**). We conclude that the LC group has considerably diminished abilities in processing speed and attentiveness, but did not show more severe cognitive deficits affecting learning and spatial memory that are seen in more severe disease models using the IntelliCage, such as in hippocampal brain lesions ^43^.

### No severe impairment of spatial learning in LC mice

Spatial memory is responsible for the recording and recovery of information needed to plan a course to a location and to recall the location of an object or the occurrence of an event ^44^. To test spatial memory, we performed place preference/place learning and place reversal tests, which were conducted for 2 days and 4 days, respectively. These tests examine basic spatial learning and memory in both groups ^45^. On day 2 from the start of the test (**Figure 1A**), each mouse was semi-randomly assigned a corner chamber where they could perform a nose-poke to open the door for 8 seconds to receive 5% sucrose water, while the other 3 corners remained inaccessible to them. On day 4 from the start of the test, place reversal started, and each mouse was assigned to a different corner. The percentage of correct visits with nose-pokes out of the total number of visits with nose pokes was calculated per day, as seen in **Figure 1E**. The overall correct percentage of the control group was 56.98% while the LC group was 56.83% (p-value=0.08, two sample t-test). This data indicates that while the control group performed slightly better than the LC group, more basic cognitive functioning was not severely impaired in the LC mice as seen using the IntelliCage in other disease models, like Alzheimer’s disease ^46^.

### Higher-order cognitive abilities and behavioral flexibility are significantly affected in LC mice

The serial reaction time /motor impulsivity test is a cognitive task that allows simultaneous assessment of several modalities, including attention, response inhibition, cognitive flexibility, and processing speed ^36,47,48^. This task includes a connected series of events that engages processes supporting the temporal organization of behavior, the formation of high-order associations, and the prediction of future events ^49,37^. Thus, this task has been used to explore the processes underlying a broad range of behaviors, including cognitive and biological principles of learning and memory. Using the IntelliCage setup, we evaluated the ability of each mouse to carefully follow specific steps to achieve their goals ^50^. During the test, all corner chambers were operated in the same manner. When a mouse performs a first nose poke during a visit, a trial was initiated and a 3 second delay would start, followed by the LED lights turning on. If the mouse performed a second nose poke after the 3 second delay, the door would open, and the trial would be considered successful. Any nose poke performed during the delay between the first nose-poke and the LED lights turning on would be considered a premature response and the trial would fail, necessitating the mouse to leave the corner chamber and re-enter to initiate another trial. If a second nose-poke was not done despite the LED lights turning on, this would be a semi-failed trial. If a trial was performed correctly on one side of the corner chamber but multiple nose pokes were performed on the other side, this would be a semi-successful trial.

When calculating the percentage of successful and semi-successful trials over the total number of trials, we found that the control group was more successful (44.41%) than the LC group (38.84%). When the number of semi-successful trials was eliminated, the control group (34.92%) still outperformed the LC group (28.93%). When the number of semi-failed trials was also eliminated, it was again found that the control group (39.44%) performed better than the LC group (31.97%) (**Figure 1F**). After eliminating the semi-successful and semi-failed trials, the percentage of successful trials per hour per day for the control group is significantly higher than the LC group (p-value= 0.004, two sample t-test). Therefore, the LC mice consistently showed more cognitive deficits in processing speed, response inhibition, and the ability to perform a series of tasks in a specific order (**Figure 1F**).

To further determine whether cognitive measurements could distinguish between LC and WT groups, we employed the unsupervised dimensionality reduction technique, Principal Component Analysis (PCA) (**Figure 1 G-I**). Each point represents a data sample, projected onto the first two principal components (PC1 and PC2). This approach helped visualize the variability of the LC samples and showed that although LC mice are divergent, the non-infected mice clustered away from them in every test. Therefore, behavior analyses using the IntelliCage detect and register all changes in behavior tests per mouse and can clearly cluster LC mice away from healthy non-infected mice.

### Machine learning-based analysis of cognitive tests

Machine learning (ML) is a specific approach within artificial intelligence (AI) focusing on enabling machines to learn from data without being explicitly programmed. ML applications can group entities that share similar features using two main approaches: classification and clustering. To better understand results from the behavior studies, ML techniques, specifically clustering, were used to identify groups of mice with similar behaviors. Importantly, no prior information about the groups was provided during the analysis, and no assumptions were made regarding the optimal number of groups or the assignment of subjects to each group. If the clustering analysis placed the LC and control mice into separate groups, this would signify that the behavior pattern and cognitive functions of LC mice have fundamentally changed. Using clustering analysis, we tested the hypothesis that LC affects the higher cognitive functions of infected mice without any human bias. To determine the optimum number of clusters, we used Silhouette, ^51^ which is a technique that measures how similar a data point is to its own cluster compared to other clusters, with values ranging from −1 to 1. A higher Silhouette score indicates well-separated and cohesive clusters.

In the patrol and patrol reversal tests, we identified 3 features that are relevant to clustering. The first parameter is the number of times a mouse went to the correct corner and did at least a single nose-poke to access water. The second parameter is the number of times a mouse went to the wrong corner and did a nose-poke. The third parameter is the number of times a mouse went to a wrong corner but realized that the light did not go on, hence deducing that the corner was not correct and did not try to drink from the corner. We calculated the sum of these parameters for each mouse for each hour of the day. This data matrix was then used for clustering analysis with the ***K-Means++*** algorithm ^52^. We then used ***sklearn*** library ^53^ to implement the clustering in Python. In the K-Means method, each cluster is represented by a ***centroid***. The algorithm initializes the centroids at random locations and iteratively adjusts them to minimize intra-cluster differences while maximizing inter-cluster differences. To mitigate the effect of randomness, we ran the algorithm 100 times, selecting the results associated with the highest score value. Applying the Silhouette approach ^51^, the optimal number of clusters was determined to be 2 (**Figure 2A**). Using K=2, the results revealed two distinct clusters: one cluster comprising all the LC mice and the other containing the control uninfected mice, as illustrated in **Figure 2A**. This demonstrates that the behavior of LC mice is distinguishable from the behavior of the non-infected mice.

**Figure 2:**
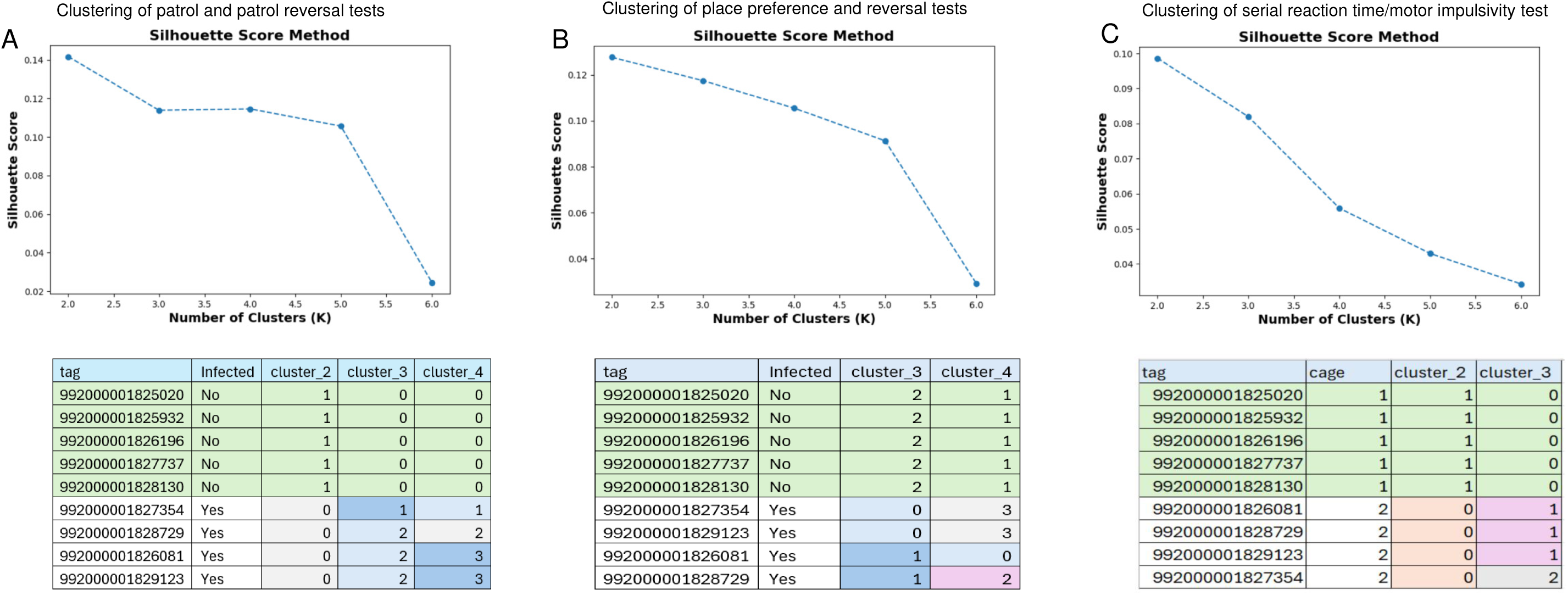
K-means clustering analysis. The range of 2 to 6 was tested to choose the number of clusters providing the highest score. **A.** The Silhouette Score of the patrol and patrol reversal test. **B.** The result of K-means analysis on the features of the mice in place preference/place learning and place reversal tests. **C.** The result of K-means analysis on the features of the mice in Serial reaction time/motor impulsivity tests. Corresponding tables below show the result of grouping the mice using 2, 3 or 4 clusters.

In the place preference and place reversal tests, parameters included the number of correct visits with nose-pokes and the number of incorrect visits with nose-pokes out of total visits with nose-pokes. Similarly, we created a matrix calculating the sum of these parameters per hour per day for each mouse. This matrix was then used by the Silhouette algorithm to determine the optimum number of clusters, which was 2, as seen in **Figure 2B**. When applying the recommended number of clusters, we found that all control mice were placed in a single cluster, while the LC mice were placed in the other clusters (**Figure 2B**). This proves again that the behavior of non-infected mice is different from their LC counterparts.

In the serial reaction time test/motor impulsivity test, the parameters included 2 types of successful trials – when a mouse successfully completes a trial on one side of the corner chamber or on both sides – and one type of semi-successful trial – when a mouse successfully completes a trial on one side of the corner chamber but fails on the other side. Similarly, two types of failed trials – when a mouse fails a trial on one side of the corner chamber or on both sides and one type of semi-failed trials – when a mouse fails to perform a second nose-poke after the LED lights go on – were used to distinguish failed trials. Similarly to the previous experiments, we calculated the total number of the different types of successes and failures for each mouse per hour per day. We created the dataset from the list of these parameters, and K-means was used to determine the optimum number of clusters. As shown in **Figure 2C**, the optimum number of clusters was determined to be 2 and after applying K=2, we obtained the classification. The LC and control mice again were assigned different clusters, providing further evidence that the cognitive function of LC mice was different from that of the control mice.

### Parallel differential expression of learning- and memory-related genes in the brains of LC mice and human patients suffering from LC-related cognitive dysfunction

We analyzed published transcriptomic data from brains of human patients who suffered from LC-related cognitive dysfunction and demonstrate the dysregulation of genes linked to cognition, learning (**Figure 3A**), synaptic functions (**Figure 3B**), and blood brain barrier pathways (BBB) (**Suppl Figure 3B**). To determine whether LC mice had similar underlying features in their transcriptomes, we isolated brains from LC mice, 30 days after initial intranasal SARS-CoV-2 infection (**Figure 1A**), and from matching uninfected controls, then performed bulk RNAseq analysis. Brains of SARS-CoV-2-infected mice show extensive divergence in gene expression patterns from uninfected control mice (**Suppl Figure 3A**). Functional analysis of differentially expressed genes in mouse brains revealed an enrichment for genes corresponding to cognition, learning and synaptic pathways **(Figure 4A-C**). To understand the mechanism underlying cognitive impairment, we analyzed genes related to inflammation, cell activation and BBB function ^54–56^. We found increased expression of neuroinflammation-related genes (**Suppl Figure 1A-F**). We also observed increased expression of microglia and astrocyte activation genes in the brains of LC mice (**Suppl Figure 1C** and **E**). Genes related to the disruption of BBB functions were also altered in brains of LC mice (**Suppl Figure 3C**), which mimics what we demonstrate in the brains of human subjects who had cognitive impairment (**Suppl Figure 3B**) ^57^. Consistent with these functional deficits in the LC mice, RNA sequencing identified differentially expressed genes associated with each of these pathways (e.g., GRIN1, GRIA2, MAPK3) compared to non-infected control mice (**Suppl Figure 3D**). Particularly the expression of the brain-derived neurotrophic factor (BDNF) is reduced in LC mice (**Figure 4B** and **Suppl Figure 1B**). Likewise, the expression of CAMK, which plays critical roles in neuronal signaling in humans, exhibited altered expression in LC mice (**Suppl Figure 3A** and **D**). Furthermore, as reported by countless human studies ^15,58–61^, we found here that the coagulation pathway was substantially altered in LC mice (**Suppl Figure 2B** and **C**) ^62–64^.

**Figure 3:**
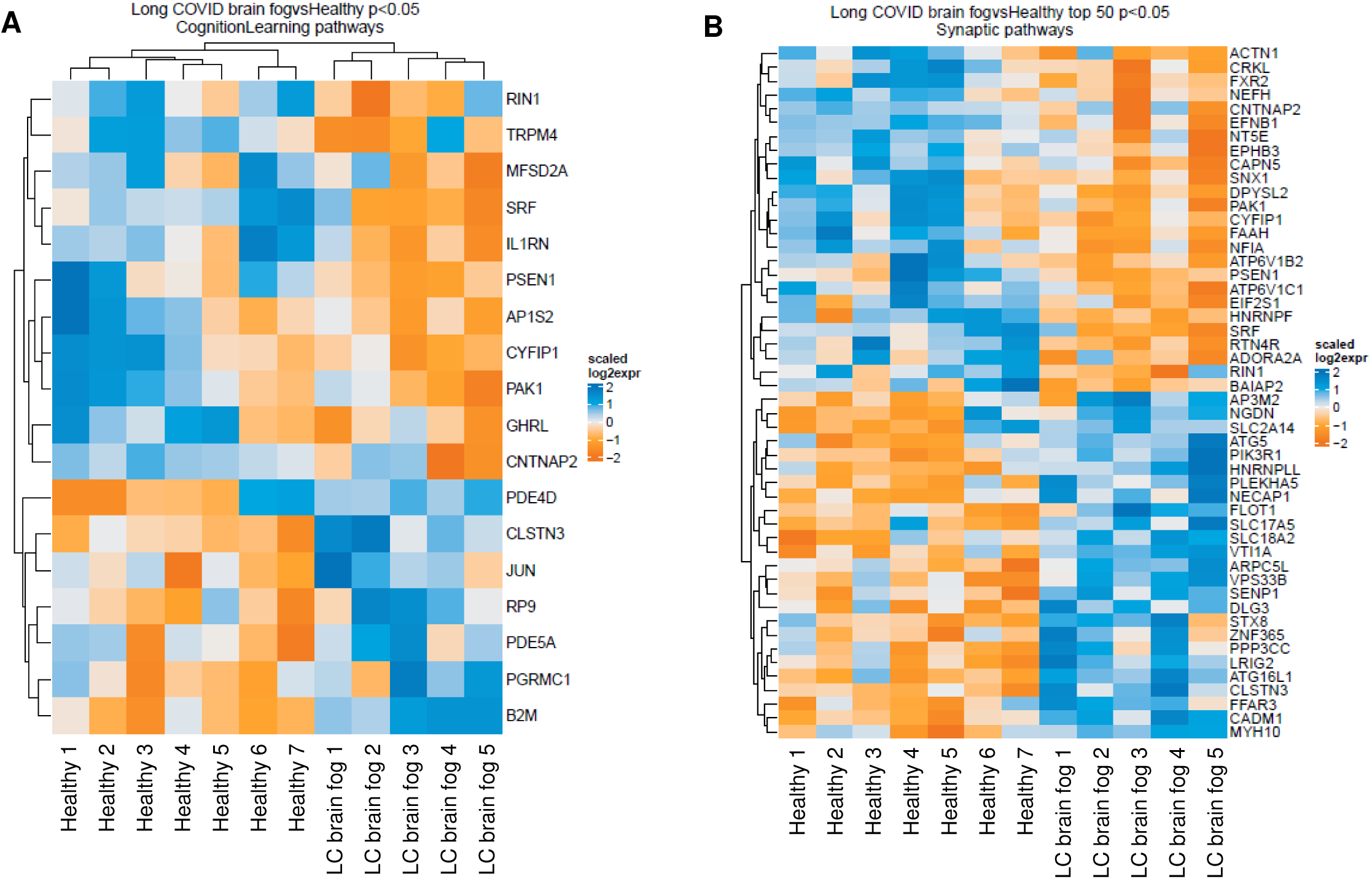
Differential expression of genes involved in cognition and learning pathways in the brains of LC individuals suffering from cognitive dysfunction. The analysis of publicly available RNA sequencing data of brain samples from subjects with LC-related cognitive dysfunction (brain fog) and healthy donors (GSE251849) revealed that the genes related to memory (**A**) and synaptic functions (**B**) are differentially expressed in individuals with LC-related cognitive dysfunction (N=5) in comparison with healthy individuals (N=7).

**Figure 4:**
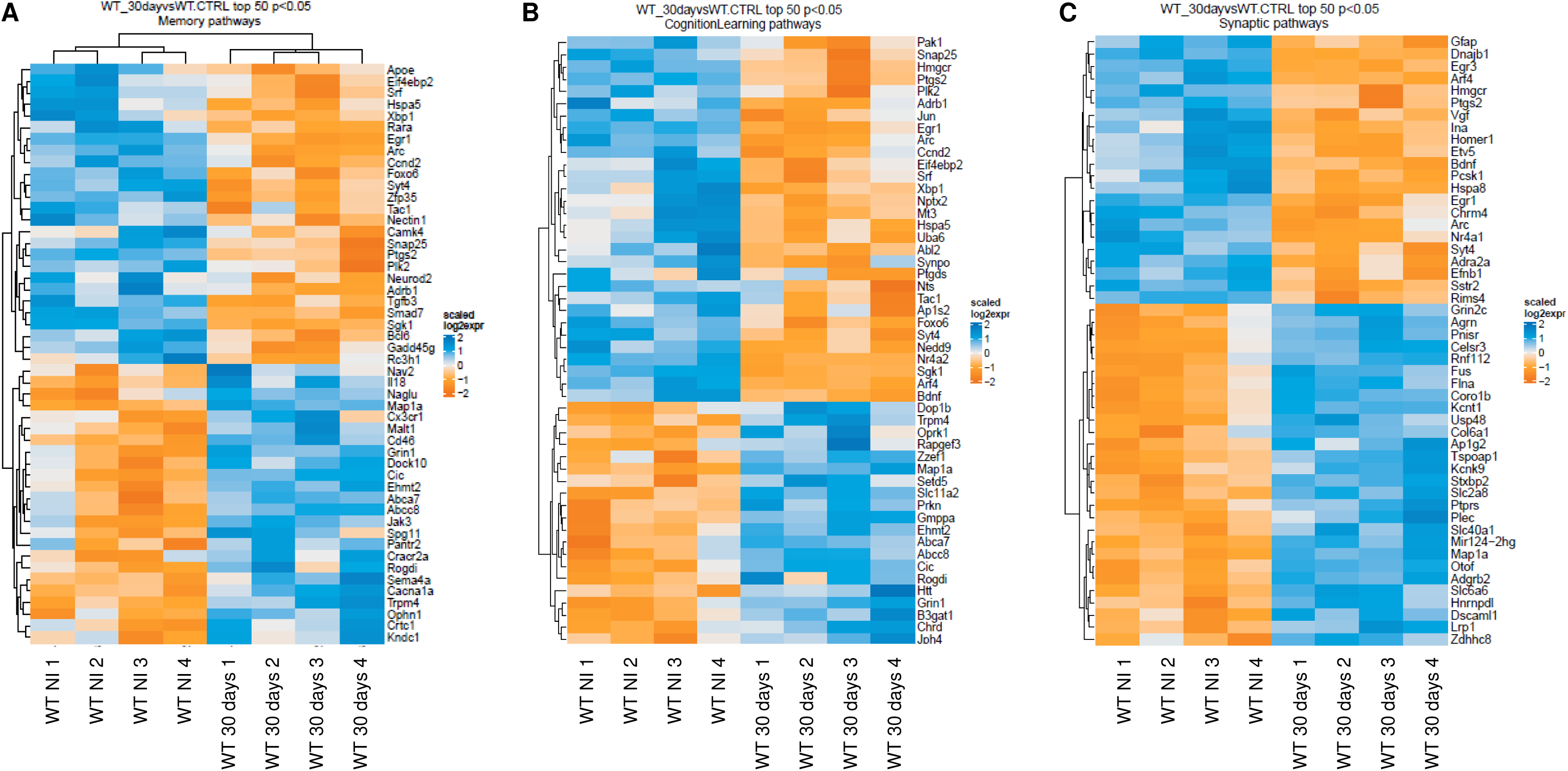
Dysregulation of genes involved in learning, cognition and synaptic functions in the brains of LC cognitive dysfunction mice. (**A-C**) Heat maps of significantly differentially expressed genes related to memory, cognition and synaptic pathways from the brains of LC mice 30 days post-infection in comparison with non-infected mice (N=4).

Then, the significant gene expression changes (p-value <0.05) were contrasted in human and mouse groups to understand how the transcriptional landscape in terms of directionality (either significantly upregulated or downregulated) changed in both species (**Suppl Figure 4A** and **B**). Functional enrichment analysis for top 10 enriched genes from our mouse data and our analysis of published human transcriptomic data of differentially expressed genes separately (**Figure 5A** and **B**) or together (**Figure 5C**) demonstrate the similarities in disrupted pathways related to cognitive dysfunction in both species. Venn diagram shows that the expression of 50 common genes was altered in the brains of mice and humans with LC-related cognitive dysfunction (**Figure 5D**). Notably, within the 50 common genes between human and mice, 29 genes were defined as significant in both groups (**Figure 5E**). Therefore, mice with cognitive dysfunction exhibit differential gene expression in pathways that mirror those altered in the brains of human subjects with LC-related cognitive dysfunction.

**Figure 5:**
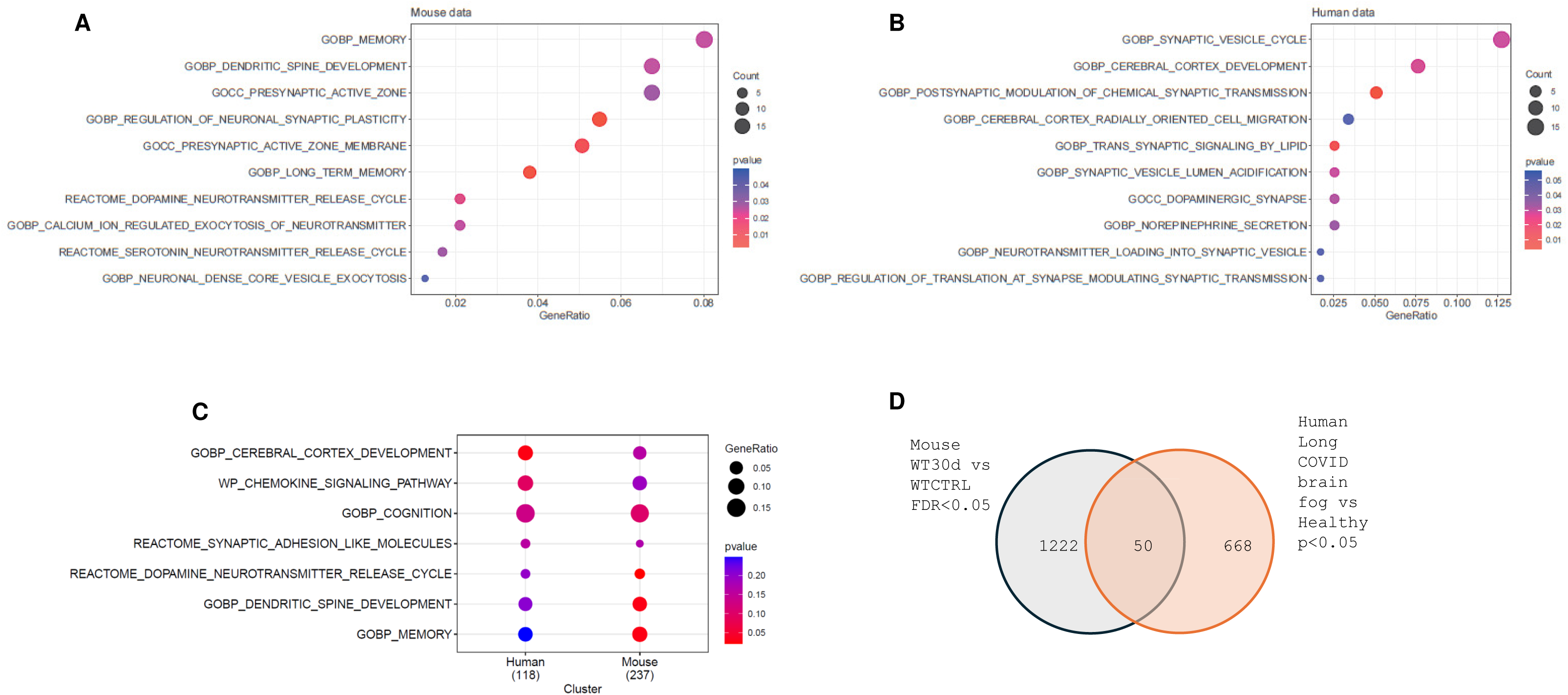
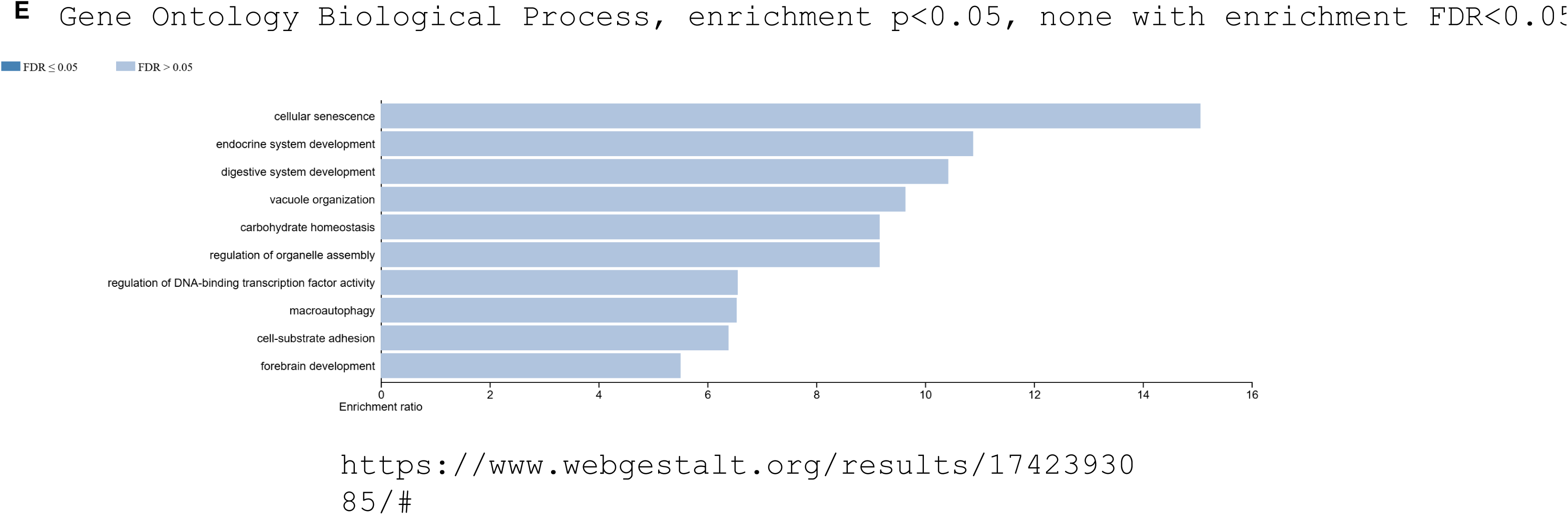
Parallel dysregulations of genes involved in similar pathways in brains of mice and humans with LC-related cognitive dysfunction. Enriched pathways related to memory, synaptic functions and neurotransmitters in mouse (**A**) and human (**B**) brains. **C**. Overlapping pathways in humans and mice with altered gene expression. **D**. The Venn diagram represents unique and shared differentially expressed transcripts in brains of LC mice (blue) and humans (red) suffering from cognitive dysfunction. In **A**, **B** and **C**, *P* values are color coded. The number of expressed genes is size coded. **E**. Bar chart enrichment of Gene Ontology Biological Processes. Gene terms enriched within the 29 genes defined as significant in both human Long COVID brain fog vs Healthy (p<0.05) and mouse WT_30d vs WT_CTRL (FDR<0.05). Human healthy controls (N=7), LC patients with brain fog (N=5). Non-infected (NI) mice N=4, LC mice (30 days post infection) N=4.

## Discussion

This is the first report to characterize complex behavioral deficits in LC mice using the IntelliCage. We demonstrate that several weeks following intranasal SARS-CoV-2 infection and in the absence of live virus, mice exhibit behavioral changes related to working memory, higher-order cognitive function and flexibility. These findings are supported by altered expression of genes involved in memory, learning, neurotransmission, neuroinflammation and cell activation in the brains of LC mice which copies transcriptional changes in patients with LC-related brain fog.

Using the IntelliCage, a small number of mice can be well characterized and distinguished, unlike the traditional behavior tests ^33,46^. Each mouse can go through several automated complex behavioral tests with no human interference. The activity of mice can also be followed throughout each test and can be repeated as frequently as needed. Machine learning was applied to analyze whether there was a distinct difference in behavior and learning between healthy non-infected and LC mice cohorts by assessing each subject’s activity level and behavior per hour, per day, and then clustering them into groups according to their behavioral pattern and cognitive abilities. Utilizing this approach, we compared the performance of LC and non-infected mice and identified a distinct behavior pattern between the groups. Notably, while the healthy mice clustered tightly together, the LC mice clustered separately and demonstrated subtle variability in their behavior (**Figure 1 G-I**). Notably, similar variable phenotypes are found in human subjects with brain fog ^65,66^. These findings, using the IntelliCage, highlight the robust analytical capabilities of the system, which effectively differentiates individual subjects despite a minimal sample size and inherent variability among mice. In humans, not every SARS-CoV-2-infected patient will report symptoms of brain fog after the resolution of infection; however, very subtle temporary effects on cognitive functions cannot be evaluated or excluded in patients recovering from SARS-CoV-2.

The IntelliCage behavioral testing platform revealed significant differences in working memory, higher cognitive abilities, and behavioral flexibility between LC and control mice. The tasks that were impaired in LC mice include patrol reverse test and serial reaction time/motor impulsivity tests. These behavioral tests assess cognitive flexibility, reversal learning, processing speed, attention, and inhibiting impulsivity, and are mediated by the prefrontal cortex ^67^. The tasks that were not significantly impaired by LC involved place learning, which assessed spatial exploration and memory functions which are mediated by the hippocampus ^33^. More tests can be designed in the IntelliCage to explore additional behavioral abnormalities. Taken together, these findings indicate that the IntelliCage system, compared with traditional paradigms, has a higher sensitivity for detecting subtle cognitive and behavioral differences that make up the complex cognitive dysfunction phenotype in LC mice.

Differential gene expression patterns in mouse models during the early stages of SARS-CoV-2 infection have been reported ^68,69^. However, here we show that even 30 days post-infection, the gene expression profile in the brain remains distinct from healthy controls. Mechanistically, neuroinflammation is one of the major drivers of cognitive dysfunction in LC and other neurological disorders ^70,71^. Altered expression levels of pro-inflammatory cytokine genes (**Suppl Figure 1F** and **2A**), particularly the chemokine CCL11 and its family members (**Suppl Figure 1D**), have been linked to cognitive impairments observed in patients with persistent symptoms ^5, 72^, and in human subjects suffering from LC-related cognitive dysfunction ^4,73,74^.

Notably, we observed increased expression of microglia and astrocyte activation which has been implicated in virus-induced cognitive impairment ^75^, memory disturbances, fatigue and insomnia ^75^. Notably, microglial activation and altered coagulation are strongly linked with neurological malfunctions in LC ^58^. Microglial activation is associated with loss of oligodendrocytes leading to myelin loss which impairs the structure and function of neuronal networks ^4,16,75^. There is evidence from post-mortem examinations of the brains of COVID-19 patients of alterations in astrocytes, microglia and myelin pathways ^76 57,77^. Reactive-astrocyte-mediated loss of the BBB integrity is also linked to brain pathology in LC patients ^57^, like what we observe in the LC mouse.

Reversal learning in LC mice was altered compared to non-infected controls, indicating impairments in higher-order cognitive functions, including cognitive flexibility, processing speed, attention, and inhibitory control. These cognitive domains are tightly linked to dysregulation in key neurotransmitter systems, particularly serotoninergic, dopaminergic, cholinergic, and glutamatergic pathways ^24,78,79^. The expression of genes involved in these pathways was significantly altered. Importantly, BDNF which is a protein involved in the regulation of neuroplasticity, neurogenesis, and neuronal differentiation, and has been linked to neurological disorders, psychiatric disorders and LC patients with brain fog ^80,81^. Since in AD patients, low levels of BDNF provokes amyloid beta (Aβ) ^82^ and Tau ^83,84^ accumulation, it is plausible that LC actually accelerates AD in LC patients as suggested by several studies by reducing BDNF (**Figure 4B**) ^85^. Likewise, the expression of genes related to synaptic plasticity, memory consolidation, and neuronal signaling such as CAMK, in humans, exhibited altered expression in the brains of LC mice as well. As reported by countless human studies ^15,58–61^, we found here that the coagulation pathway was substantially altered in LC mice. Together, our data demonstrate that the LC-related cognitive dysfunctions are recapitulated in the mouse several weeks after SARS-CoV-2 infection while providing a mechanistic insight underlying the behavioral changes observed in LC mouse model in comparison with patients with brain fog.

In conclusion, our data establishes the IntelliCage as a valuable platform for studying LC-associated cognitive disfunction in mice and could be used to identify new biomarkers, drug targets and for screening potential pharmacological interventions.

---

**Supplementary Figure 1.**
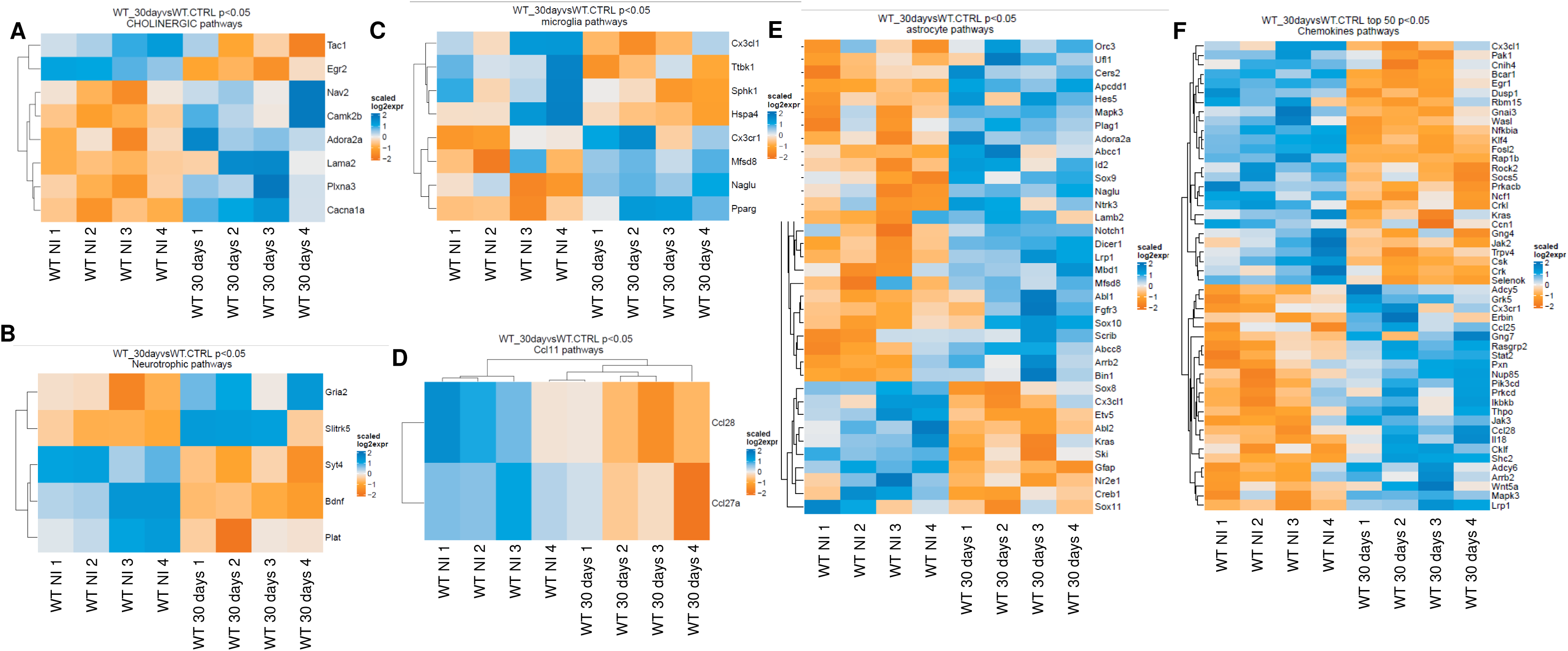
**A-E** Heat maps for altered gene expression involved in chemokines pathways, CCL11, microglia and astrocyte activation.

**Supplementary Figure 2.**
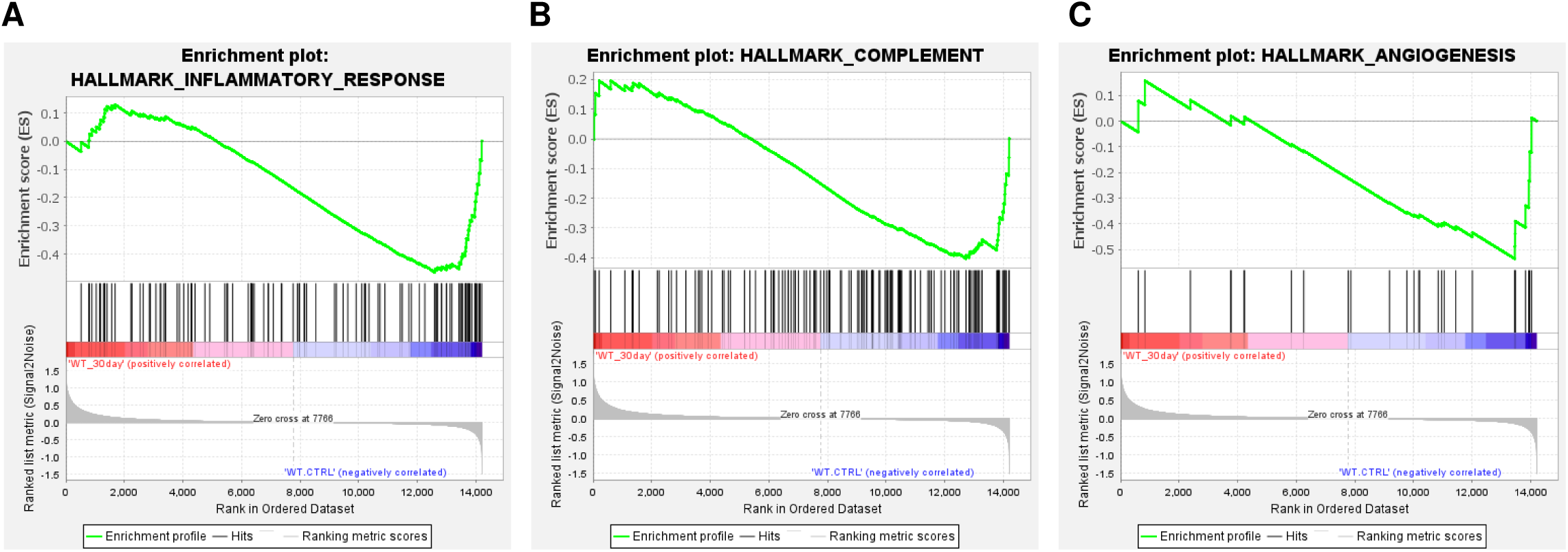
Gene set enrichment analysis results of the difference between mice 30 days post SARS-CoV-2 infection and non-infected mice for inflammatory (**A**), complement (**B**) and angiogenesis (**C**) pathways.

**Supplementary Figure 3.**
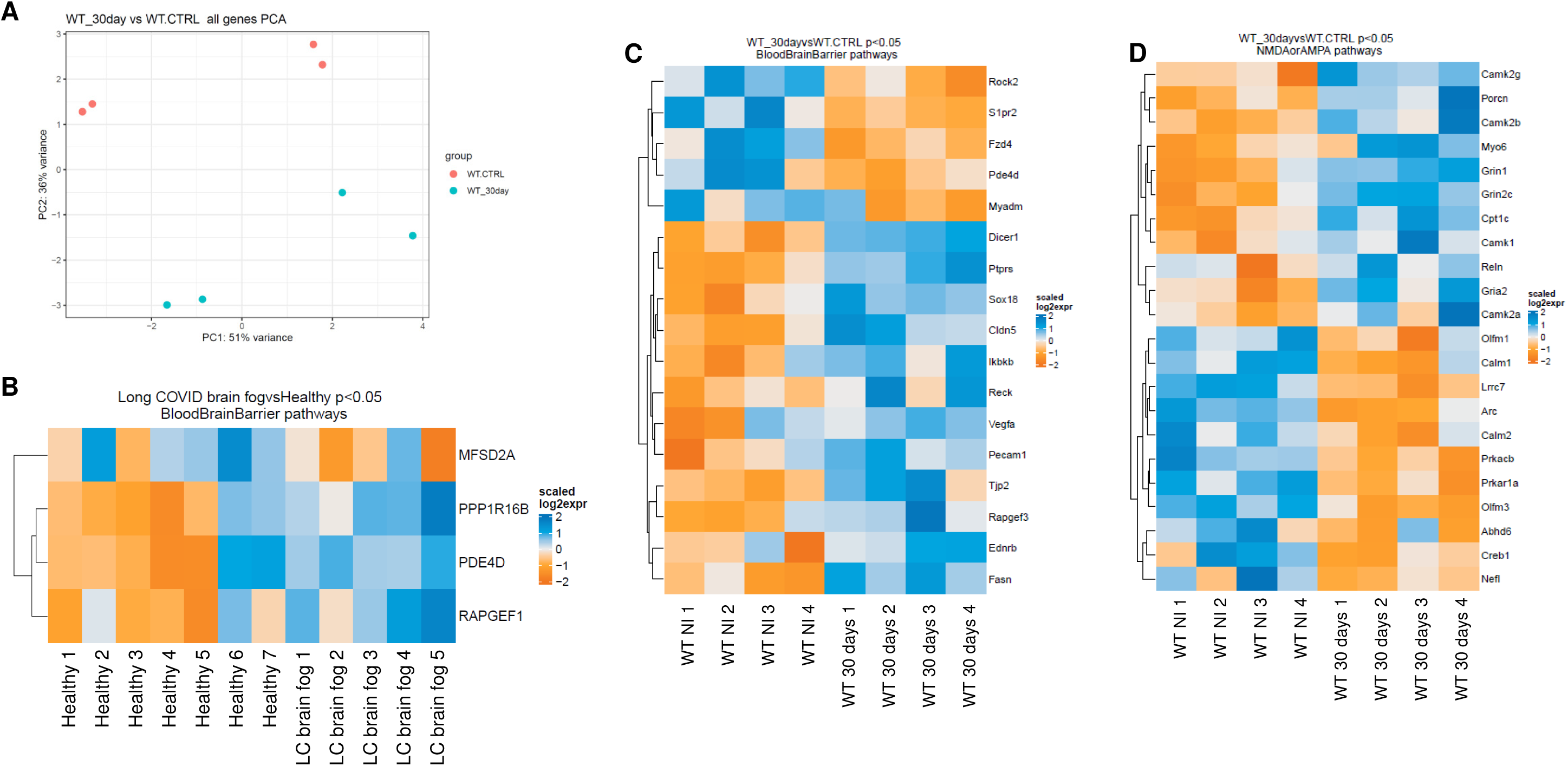
**A.** PCA plot based on the gene expression profiles in LC mice versus control uninfected counterparts. (**B**, **C** and **D**) Heat maps for altered gene expression involved in blood brain barrier function in human patients with LC-related brain fog (**B**), and LC mice compared to non-infected controls (**C**) and human patients with LC compared to controls (**C**).

**Supplementary Figure 4.**
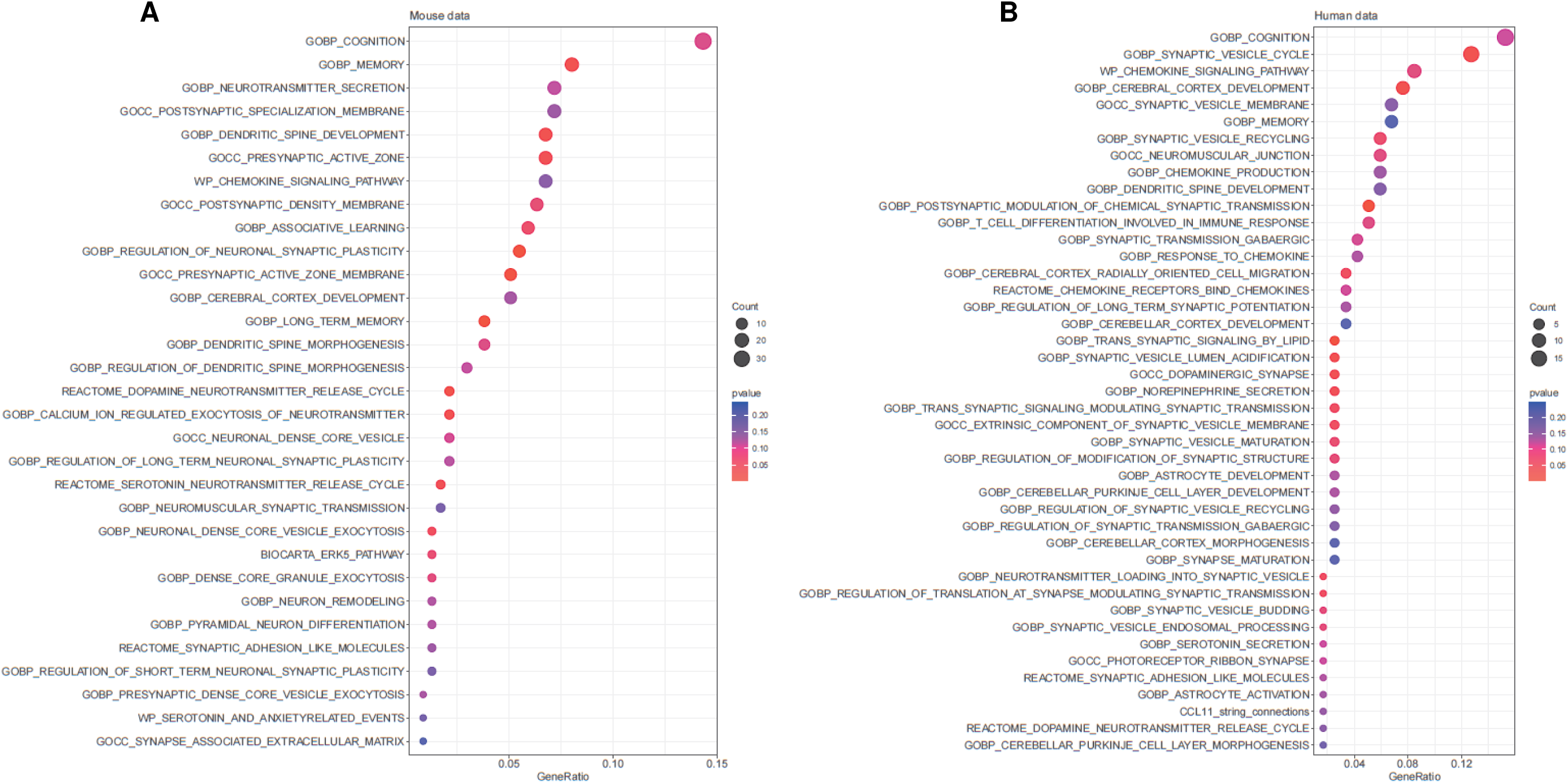
Dot plot of enriched pathways in mice and human patients with LC-related cognitive dysfunction. Shading coded by p value of enrichment and size coded by number of differentially expressed genes in gene term. The X axis location determined by ratio of differentially expressed genes to total number of genes in gene term.

## Methods

### Biosafety

All experiments with live SARS-CoV-2 were performed in the OSU BSL3 biocontainment facility. All procedures were approved by the OSU BSL3 Operations/Advisory Group, the OSU Institutional Biosafety Officer, and the OSU Institutional Biosafety Committee.

### Viruses and titers

Mouse adapted SARS-CoV-2, variant strain MA10^86^, generated by the laboratory of Dr. Ralph Baric (University of North Carolina) was provided by BEI Resources (Cat # NR-55329). SARS-CoV-2 strain USA-WA1/2020 was also provided by BEI Resources (Cat # NR-52281). Viral stocks from BEI Resources were plaque purified on Vero E6 cells to identify plaques lacking mutations in the polybasic cleavage site of the Spike protein via sequencing. Non-mutated clones were propagated on Vero E6 cells. Virus aliquots were flash frozen in liquid nitrogen and stored at −80 C. Virus stocks were sequenced to confirm a lack of tissue culture adaptation in the polybasic cleavage site. Virus stocks and tissue homogenates were tittered on Vero E6 cells as previously described ^60^.

### Mice

C57BL/6 wild-type (WT) mice aged 11-12 weeks were obtained from the Jackson Laboratory (Bar Harbor, ME, USA). All infections were performed intranasally on anesthetized mice with viruses diluted in sterile saline. All mice were housed together in a pathogen-free facility for at least 2 weeks ^60^. Experiments were conducted with approval from the Animal Care and Use Committee at the Ohio State University (Columbus, OH, USA) which is accredited by AAALAC International according to guidelines of the Public Health Service as issued in the Guide for the Care and Use of Laboratory Animals.

### The IntelliCage

The IntelliCage apparatus (39 × 58 × 21 cm) has four chambers in each corner accessible through an open tunnel. Within each corner chamber, there are two doors (left and right) that control access to the water bottles and above each door are 3 LED lights that can be turned on or off depending on the program running. A radiofrequency identification transponder (Standard Microchip T-VA, DataMars, Lamone, Switzerland; and Trovan, Melton, UK) was implanted into the dorso-cervical region of mice under isoflurane inhalation anesthesia in order to track each mouse in the corner chambers. Each RFID is detected by the ring antenna in each corner. The condition of light was as follows: the light period, 06:00–18:00 local time, and the dark period, 18:00–06:00 (Light–Dark 12:12 h). The order of the test battery is the following: free adaptation (3 days), nose-poke adaptation (2 days), patrolling test training (1 day), patrol test (3 days), patrol reversal test (3 days), place preference/learning test (2 days), place preference reversal test (4 days) and serial reaction time/motor impulsivity test (7 days). Four LC and five uninfected age-matched female mice were used and analyzed in the IntelliCage. The IntelliCage tests were initiated 55 days post-SARS-CoV-2 infection. The home cage-like conditions in the IntelliCage provide a unique experimental opportunity to assess activity freely initiated by animals. Additionally, short- and long-term habituation and circadian rhythm can be recorded. Living in groups does not affect spontaneous activity of individual mice in the IntelliCage ^33^.

### IntelliCage adaptation phase

Mice were introduced within the IntelliCage, eight weeks post SARS-CoV-2 infection. There was a 3-day free adaptation period ^35^ in which all the doors in all the corner chambers remained open to allow the mice to acclimate to their new environment and to ensure all mice were able to access the corner chambers and drink the tap water provided. This was followed by a 2-day nose-poke adaptation period ^87^. During this phase, all the doors in all the corner chambers were closed and mice had to learn that they must perform a nose poke to open the door for 10 seconds. After the 10 seconds have passed, the door would close again and the mouse would have to leave the corner chamber, re-enter that chamber (or enter another one) and perform another nose poke at the closed door in order to drink again. As all the mice successfully learned this process, the actual testing period commenced.

### IntelliCage patrol and patrol reversal test

The patrol test ^88,89^ and patrol reversal test ^87^ were conducted over a total of 7 days (1 day training and 3 days each for the patrol and patrol reversal tests). During the training period, mice were allowed access to all corners of the Intellicage and would need to nose poke to open doors and access water. Throughout any drinking periods, the LED lights would turn on and would teach the mice to associate the LED light being on with access to water. During the testing period, the mice had to learn that there would only be one working corner chamber at a time (assigned semi-randomly to each mouse) that they could perform a nose poke at to open the door. When in the correct corner, the lights above the doors would turn on, thus acting as a positive cue they would need to process. After visiting a correct corner, the working corner chamber would change in a clockwise (in the patrol test) and then a counterclockwise orientation (in the patrol reversal test). Therefore, the mice had to learn the pattern that the corners were changing in, remember the previously correct corner and then plan prospectively to reach the newly correct corner. Furthermore, mice had to also actively process in the moment that the lights going on meant they were in the correct corner (and that they may proceed to perform a nose poke to get to drink) and that the lights not turning on meant they were in the incorrect corner (and thus deduce that performing a nose poke would be fruitless and leave). As a result, 3 parameters were assessed and calculated throughout this experiment: the number of times a mouse entered the correct corner chamber and performed at least one nose poke out of the total number of visits with nose pokes, the number of times a mouse entered an incorrect corner and performed at least one nose poke out of the total number of visits with nose pokes and the number of times a mouse entered an incorrect corner but did not perform a nose poke when the lights did not go on out of the total number of times a mouse entered an incorrect corner ^34,36,90^.

### IntelliCage place preference and place reversal test

The place preference and place reversal tests ^91^ were conducted over a period of 2 days and 4 days, respectively, using 5% sucrose water. In the place preference test, only one corner chamber offering sucrose-water would open if a mouse performed a nose poke while the other 3 would remain inaccessible to that mouse. The corner assignment was semi-randomized for each mouse and on day 4 of this experiment, the place preference reversal test began with the correct corner for every mouse changing. Thus, the mice had to learn and remember which corner was made available to them during the place preference test and then relearn and remember the new corner assigned to them during the place preference reversal test. The parameters evaluated included the number of correct visits with nose pokes out of total visits with nose pokes and the number of incorrect visits with nose pokes out of total visits with nose pokes.

### IntelliCage serial reaction test

The serial reaction time test/motor impulsivity test ^46,92^ took place over 7 days. All the corners in this test were set as working and the mice had to learn a specific series of actions to be able to get the door to open. First, a mouse would have to perform a nose-poke to initiate a trial. Then, the mouse had to wait for a 3 second delay period without doing any further nose pokes (otherwise the trial would fail) before a light went on above the door. After the lights go on, if the mouse did a second nose-poke, the door would open, and the trial would be considered successful. A trial was considered as a semi-failure if the mouse did not perform a second nose-poke and was considered a semi-success if the mouse successfully opened the door on one side of the corner chamber but performed multiple nose-pokes on the other side. Therefore, the mouse had to learn how to follow a series of procedural steps and actively pay attention to the correct order and cues in order to succeed. The parameters calculated during the test included the number of trials initiated, the number of trials failed during the delay period, the number of semi-failed trials, the number of semi-successful trials and the number of completely successful trials.

### RNAseq and data analysis

As we previously described ^60^, total RNA was extracted from the brains of wild-type (WT) mice 30 days post SARS-CoV-2 infection and from sex- and aged-matched non-infected controls by TRIzol reagent (Thermo Fisher Scientific, 15596026) according to the manufacturer’s instructions. RNA cleaning and concentration were done using Zymo Research, RNA Clean & Concentrator-5 kit (cat# R1015) following the manufacturer’s protocol. Fluorometric quantification of RNA and RNA integrity analysis were carried out using RNA Biochip and Qubit RNA Fluorescence Dye (Invitrogen). cDNA libraries were generated using NEBNext Ultra II Directional (stranded) RNA Library Prep Kit for Illumina (NEB #E7760L). Ribosomal RNA was removed using NEBNext rRNA Depletion Kit (human, mouse, rat) (E #E6310X). Libraries were indexed using NEBNext Multiplex Oligos for Illumina Unique Dual Index Primer Pairs (NEB #644OS/L). Library prep generated cDNA was quantified and analyzed using Agilent DNA chip and Qubit DNA dye. Ribo-depleted total transcriptome libraries were sequenced on an Illumina NovaSeq SP flow cell (paired-end 155bp format; 35-40 million clusters, equivalent to 70-80 million reads. Library preparation, QC, and sequencing was carried out at Genewiz.

Sequencing data processing and analysis were performed by the Bioinformatics Shared Resource Group (BISR) at the Ohio State University using previously published pipelines ^93^. Briefly, raw RNAseq data (fastq) were aligned to mouse reference genome (GRCm38) using hisat2 (v2.1.0) ^94^ and converted to counts using the ‘subread’ package (v1.5.1) ^95^. Low expressed counts were excluded if more than half of the samples did not meet the inclusion criteria (2 CPM). Data were normalized using ‘voom’ and statistical analysis for differential expression was performed with ‘limma’ ^96^. For data visualization, DESeq2 rlog transformation was used for principal component analysis (PCA). Volcano plots were generated with ‘EnhancedVolcano’ and heatmaps were generated ‘ComplexHeatmap’ using R. Over-representation enrichment analysis was performed with ‘clusterProfiler’ ^93^, and EnrichR for enriched pathways and regulatory networks ^97^.

### Statistical analysis

Data was analyzed and comparisons between the groups were done using java apache math3 library version 3.6.1 to calculate t-test scores. Comparisons between groups were conducted with either two-sample t-test or ANOVA followed by Tukey’s multiple comparisons test. Adjusted P<0.05 was considered statistically significant and P<0.1 was considered marginally significant.

### RNAseq Data availability

RNA sequencing data comparing LC WT and non-infected WT mice brains are shared through Gene Expression Omnibus with accession number GSE292780.

## Author Contributions

Conceptualization, A.O.A., H.M.A., M.M.S., M.E., O.K-C., R.M.B. Experiments and data acquisition, H.M.A., M.M.S., M.E., S.F., R.E., M.C., R.P., O.W.; Generation of critical reagents, R.E., G.G., J.O., J.Y., D.B., E.C-B., P.B., J.L., M.E.P., S.S., E.O., M.K.C.; Data Analysis, A.W., X.Z., M.P., M.A.; Writing – Original draft, H.M.A., M.M.S., M.A., R.M.B., A.O.A.; Writing – Review and editing, all authors.; Project Administration, A.O.A.; Supervision, A.O.A., E.C.-B., P.N.B., E.O.; Funding Acquisition, A.O.A., E.C-B., P.N.B., J.L., X.Z, E.O.

## Acknowledgements

A.O.A., E. C-B, P.N.B, J.L, X.Z. are supported by NIH grant P01 AI175399. A.O.A, S.S. are supported by R01 AI154553, R01 AI157205. A.O.A and J.Y. are supported by R01 HL168501. A.O.A is supported by R01 AG082113. P.N.B. is supported by R01AI1145144. E.O is supported by U54CA260582. M.P. is supported by R01 AI093848, U19 AI42733, CFF MCCOY19R0.

